# Controlling taxa abundance improves metatranscriptomics differential analysis

**DOI:** 10.1101/2022.03.30.486415

**Authors:** Zhicheng Ji, Li Ma

## Abstract

A common task in analyzing metatranscriptomics data is to identify microbial metabolic pathways with differential RNA abundances across multiple sample groups. With information from paired metagenomics data, current differential methods control for either DNA or taxa abundances to address their strong correlation with RNA abundance. We discovered that when either DNA or taxa abundance is controlled for, RNA abundance still has a strong partial correlation with the other factor. Thus, both factors need to be controlled for in the differential model. In both simulation studies and a real data analysis, we demonstrated that controlling for both DNA and taxa abundances leads to superior performance compared to only controlling for one factor.

## Main

Metatranscriptomics [1] and metagenomics [2] studies by shotgun sequencing profile taxonomic abundances and RNA and DNA abundances of genes and gene pathways in a microbial community. They provide a more comprehensive landscape of microbial functions and activities than traditional technologies such as 16S rRNA sequencing [3] that only profile taxonomic abundances. Metatranscriptomics and metagenomics profiles of hundreds of individuals have been generated in large consortium studies such as The Integrative Human Microbiome Project (iHMP) [4] and The Inflammatory Bowel Disease Multi’omics Database (IBDMDB) [5]. A fundamental goal in analyzing such data is to understand how changes in microbial compositions or gene activities are associated with disease status, thus providing new insights into the disease mechanisms. This involves the comparisons of RNA, DNA, and taxa abundances across samples and the identification of pathways with differential RNA abundances between sample groups.

While significant progress has been made to develop methods for differential compositional analysis [6], methods to identify differentially expressed (DE) gene pathways for metatranscriptomics data are still underdeveloped. DE analysis for metatranscriptomics data is different from that for single-organism RNA-seq data in that the RNA abundance is highly related to the taxonomic abundances and genes’ copy numbers in a microbial community with many organisms. If DE analysis is directly performed without addressing these effects, it is hard to differentiate if the DE is due to the changes in DNA or taxonomic abundances or due to the changes in relative RNA abundances independent of DNA or taxonomic changes. Thus, methods designed for single-organism RNA-seq data such as limma [7], DESeq2 [8], and edgeR [9] cannot be directly applied. To address this issue, Biobakery 3 [10] takes advantage of the paired metagenomics information measured in the same individuals, and include DNA abundance as a covariate in a linear-mixed effect model to address the strong correlation between DNA and RNA abundance. A recent study [11] benchmarks the performance of six DE models controlling for either DNA or taxonomic abundances, and concludes that the model that only controls for DNA abundance has the preferred performance.

The existing methods assume that including either DNA or taxonomic abundance in DE is sufficient to control for all confounding effects. Surprisingly, we found that if only DNA or taxonomic abundance is controlled for, there is still a strong partial correlation between RNA abundance with the other factor, meaning that the existing methods do not fully control for the confounding effects. For example, we downloaded and processed paired metatranscriptomics and metagenomics data from IBDMDB. For each feature (gene pathway in a species), we calculated the partial correlation between RNA abundances and taxonomic abundances after controlling for DNA abundances (Figure 1A-1C). There are 9.1% of all features with partial correlations < −0.3 and 11.4% of all features with partial correlations > 0.3. Likewise, we calculated the partial correlation between RNA abundances and DNA abundances after controlling for taxonomic abundances (Figure 1D-1F). There are 2.0% of all features with partial correlations < −0.3 and 43.2% of all features with partial correlations > 0.3. In both cases, we observe a large number of features with strong positive or negative partial correlations, suggesting that both DNA and taxonomic abundances need to be adjusted in DE to fully control for confounding effects.

**Figure 1.**
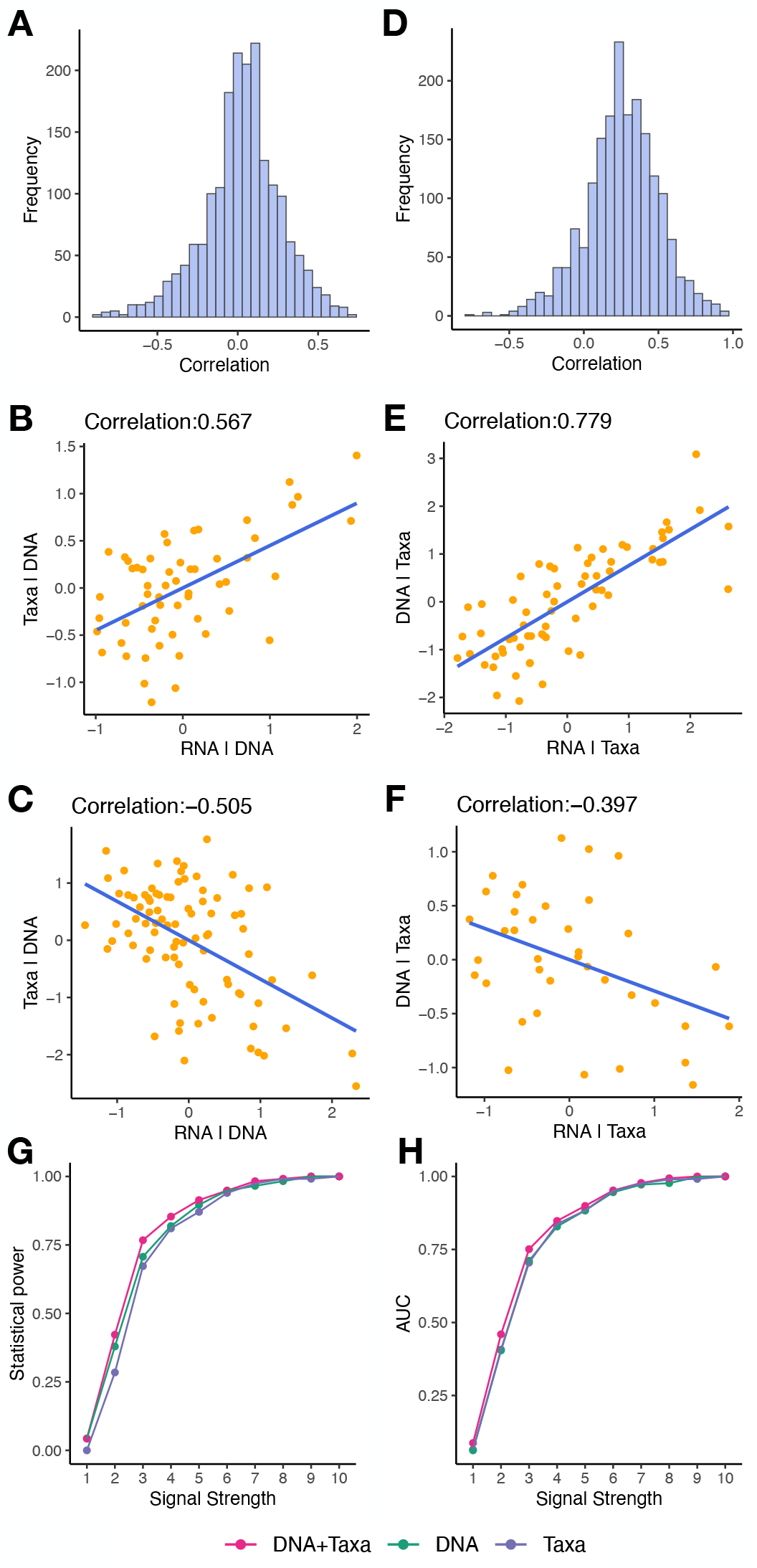
Motivation and simulation studies. **A** Distribution of partial correlations between RNA and taxonomic abundances controlling for DNA abundances. **B-C** Example features with strong positive (B) or negative (C) partial correlations between RNA and taxonomic abundances controlling for DNA abundances. x-axis shows the RNA abundances regressing out DNA abundances. y-axis shows the taxonomic abundances regressing out DNA abundances. **D** Distribution of partial correlations between RNA and DNA abundances controlling for taxonomic abundances. **E-F** Example features with strong positive (E) or negative (F) partial correlations between RNA and DNA abundances controlling for taxonomic abundances. x-axis shows the RNA abundances regressing out taxonomic abundances. y-axis shows the DNA abundances regressing out taxonomic abundances. **G-H** Results of simulation studies. x-axis shows different levels of signal strength. y-axis: statistical power at 0.05 FDR (G) or area under FDR-sensitivity curve (H).

We conducted a simulation study to systematically evaluate the performance of DE models when both or only one factor is controlled for. The simulation dataset was constructed using real RNA, DNA, and taxonomic abundance information from IBDMDB with increasing strengths of differential signals (Methods). Since the data has a longitudinal design, linear-mixed effect models were used for DE analysis (Methods). Three different models were applied for DE analysis: controlling for both DNA and taxonomic abundances (DNA+Taxa), controlling only for DNA abundances (DNA), and controlling only for taxonomic abundances (Taxa). We evaluated the performance of three models by statistical power at 0.05 FDR (Figure 1G), and area under FDR-sensitivity curve (AUC, Figure 1H). The model that controls for both factors (DNA+Taxa) has consistently the best performance in all scenarios.

As a real data example, we applied the three DE models (Methods) to the IBDMDB dataset to identify differential features between active (dysbiotic) and inactive (non-dysbiotic) time points in patients with Crohn’s disease (CD). We first performed a reproducibility analysis where all samples were randomly split into two groups and the three DE models were applied to each sample group. We evaluated how many top differential features can be reproducibly identified in both sample groups. Results identified by the model controlling for both factors (DNA+Taxa) are consistently more reproducible compared to results from other two models (Figure 2A). Since the real biological signal is more likely to reproduce itself across multiple sample groups, this result suggests that the DNA+Taxa model is more capable of identifying real biological signals. We then applied the three DE models to the full dataset. Since a rigorous gold standard is lacking, we consulted PubMed and recorded numbers of publications supporting the differential features identified by each DE model (Supplementary Table 1, Methods). DNA+Taxa model identifies 12 differential features that cannot be identified by DNA model, and 8 (67%) of them has > 30 PubMed supports (Figure 1B). DNA+Taxa model also identifies 24 differential features that cannot be identified by Taxa model, and 13 (54%) of them has > 30 PubMed supports (Figure 1B). In comparison, DNA model identifies 6 differential features that cannot be identified by DNA+Taxa model, and 1 (17%) of them has > 30 PubMed supports (Figure 1B). Taxa model identifies 4 differential features that cannot be identified by DNA+Taxa model, and 1 (25%) of them has > 30 PubMed supports (Figure 1B). These results suggest that DNA+Taxa model is able to identify more differential features, and those new features are better supported by existing literature, suggesting that results by DNA+Taxa are more biologically relevant. Figure 1C shows the names and differential status of features that are identified as differential in at least one but not all DE models. Figure 1D and 1E show two example features which are identified as significantly differential by DNA+Taxa model but non-significant by DNA or Taxa model. In both cases, RNA abundances controlled for both DNA and taxonomic abundances show much stronger differences between two sample groups compared to those when only one factor is controlled for.

**Figure 2.**
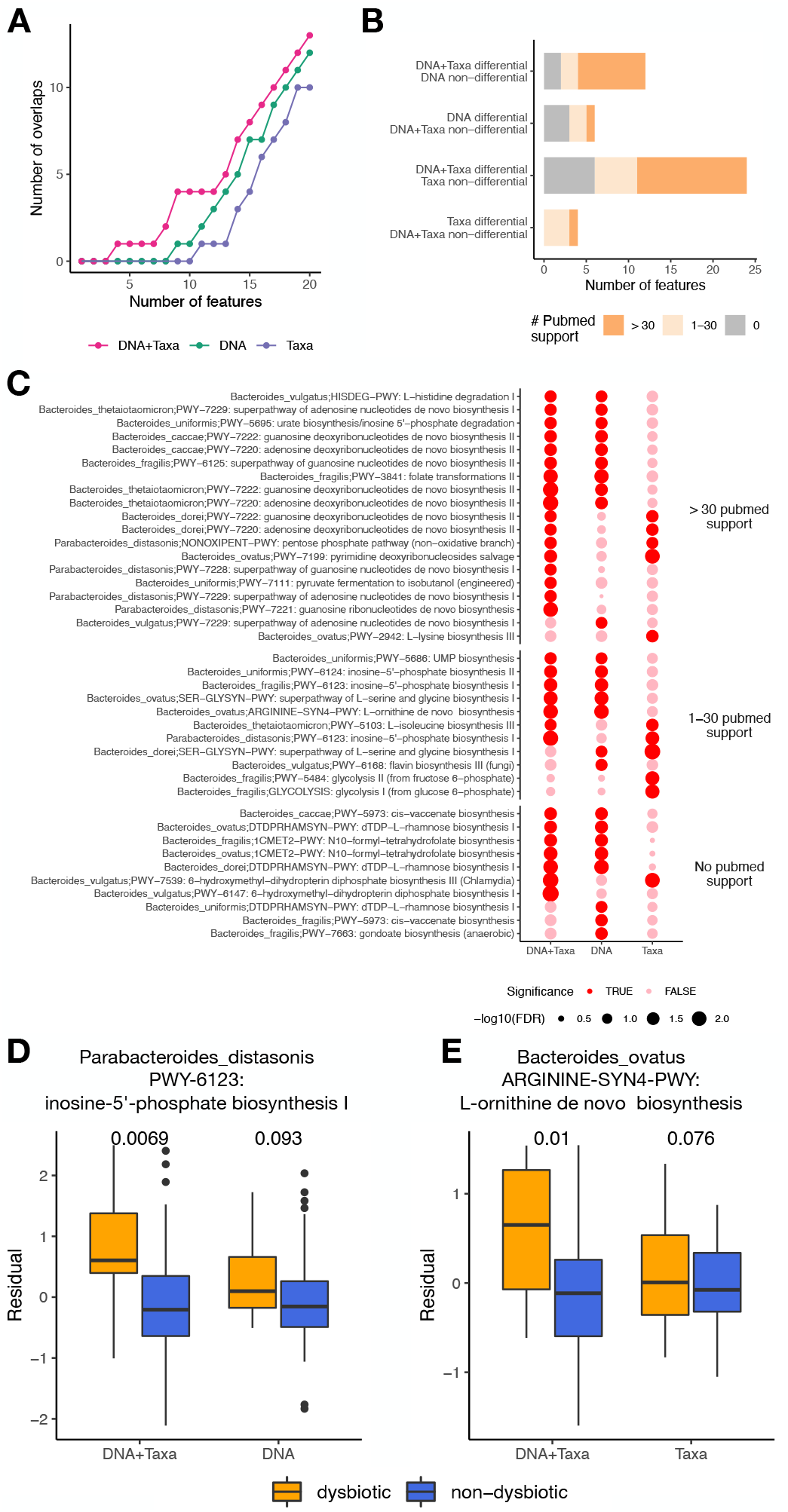
IBDMDB data analysis. **A** Reproducibility analysis. x-axis shows the number of top differential features. y-axis shows the number of top differential features identified in both sample groups. **B** Comparison of number of differential features identified by three DE models. Colors indicate the number of PubMed search hits for each differential feature. **C** Full list of differential features identified by at least one but not all DE models. Features with different levels of PubMed literature support are placed in three groups. **D** Example feature identified as differential by DNA+Taxa model but not by DNA model. y-axis shows the RNA abundances regressing out DNA and Taxa (left) or DNA (right), as well as covariates of age and antibiotic status. FDRs of statistical tests are indicated on the top. **E** Example feature identified as differential by DNA+Taxa model but not by Taxa model. y-axis shows the RNA abundances regressing out DNA and Taxa (left) or Taxa (right), as well as covariates of age and antibiotic status. FDRs of statistical tests are indicated on the top.

In summary, we reported significant partial correlations between RNA abundances and DNA or taxonomic abundance when the other factor is controlled for. We demonstrated that the DE model controlling for both DNA and taxonomic abundances outperforms DE models that only control for one factor in both simulation studies and real data analysis. This finding will provide new insights for performing DE analysis or designing new DE methods for metatranscriptomics data.

## Methods

### IBDMDB data processing

RNA and DNA abundances of microbial metabolic pathways for each species, taxonomic abundances, and sample metadata of IBDMDB dataset were downloaded from the IBDMDB website (https://ibdmdb.org/). Unmapped reads or unknown taxa are removed from the data. Downloaded RNA and DNA abundances are in cpm (counts per million), and they are further log2 transformed after adding a pseudocount of 1. Taxa abundance values are center-log transformed.

For each feature (gene pathway in a species), only samples that have all positive values in RNA, DNA, and taxonomic abundances are kept. Such filtering addresses the excessive amounts of zeroes in RNA abundances [10], and similar approach has been shown to have superior performance [11].

### Statistical Models

The IBDMDB study provides longitudinal measurements for each subject. After data filtering, we observed in some features that most subjects only have one longitudinal observation, while in other features many subjects have multiple longitudinal observations. Thus we first used a model selection procedure to determine if a linear model with only fixed effects or a linear-mixed effect model is needed for each feature. The two models differ in that the linear-mixed effect model includes a random effect indicating which subject each longitudinal observation was collected. For each feature we fitted both models and compared their model fittings using a likelihood ratio test. The p-values were obtained with asymptotic chi-squared tests and adjusted by BH procedure [12] to obtain FDRs. For all features with FDR < 0.05, linear-mixed effect models were used. For all features with FDR > 0.05, linear models with only fixed effects were used.

For simulation studies, the DNA+Taxa model is: *RNA* ~ *DNA* + *Taxa* + *group* + (1|*sub ject*). The DNA model is: *RNA* ~ *DNA* + *group* + (1|*sub ject*). The Taxa model is: *RNA* ~ *Taxa* + *group* + (1|*sub ject*). Here *RNA, DNA*, and *Taxa* represent the processed RNA, DNA, and taxonomic abundances. *group* indicates the two sample groups being compared with, and *sub ject* indicates the subjects from which samples are longitudinally collected.

For real data analysis of IBDMDB data, the DNA+Taxa model is: *RNA* ~ *DNA* + *Taxa* + *active* + *age* + *antibiotics* + (1|*sub ject*). The DNA model is: *RNA* ~ *DNA* + *active* + *age* + *antibiotics* + (1|*sub ject*). The Taxa model is: *RNA* ~ *Taxa* + *active* + *age* + *antibiotics* + (1|*sub ject*). Here *active* indicates the dysbiotic and non-dysbiotic sample groups being compared with, and *age* and *antibiotics* are the age and antibiotics status for each subject, similar to a previous study [10].

DE analysis was performed only in features with at least 10 observations in both sample groups after data filtering. All linear-mixed effect models were fitted using the lmerTest::lmer() function in R. To test for the fixed effect of interest, the Satterthwaite method [13] was used to estimate degrees of freedoms and p-values were obtained with t-tests. FDRs were obtained using BH procedure [12].

### Simulation studies

Simulation datasets were created using the processed IBDMDB data. All samples were randomly assigned to two groups with equal chances to create a null situation where no differential feature is expected between the two groups. Then, artificial differential signals were added to selected features with absolute values of partial correlations (either between RNA and DNA abundances or between RNA and taxonomic abundances) larger than 0.3. Specifically, all positive RNA abundance values across all samples and features are sorted and assigned to 10 groups, so that the minimum value in the *k*th group is no smaller than the maximum value in the (*k* − 1)th group. To add differential signals to one feature, RNA abundance values in one randomly chosen sample group are added by values randomly selected from the *k*th group. Differential features are then identified by each DE model and compared to the gold standard differential features. The simulation study is repeated for each choice of *k* (*k* = 1, 2, …, 10).

### PubMed support search for real data analysis

For each feature, we identified a keyword of its associated metabolic gene pathway. We searched PubMed with the combination of “crohn’s disease” and the keyword. For example, the PubMed search is “(crohn’s disease) AND adenosine” if the keyword is “adenosine”. We then recorded the number of publications found by PubMed.

## Availability of data and materials

The processed IBDMDB data was downloaded from the IBDMDB website https://ibdmdb.org/.

## Authors’ contributions

Z.J. and L.M. conceived the study. Z.J. performed the data analysis. Z.J. and L.M. wrote and approved the manuscript.

## Additional Files

**Supplementary Table 1 — list of differential features and PubMed literature support**

1st column: name of the feature; 2nd column: name of the gene pathway; 3rd column: keyword of the pathway used to perform literature search; 4th column: Number of PubMed hits searching for both crohn’s disease and the keyword in the third column. 5th-7th columns: If the feature is identified as differential in DNA+Taxa, DNA, or Taxa model.

